## Supplemental Display Items for "A Spatiotemporal Reconstruction of the *C. elegans* Pharyngeal Cuticle Reveals a Structure Rich in Phase-Separating Proteins"

| | Trivial Name[a] | Gene Class[b] | Name Status[c] | Systematic Name | Time of Expression[d] (hour, degrees) | Fold Enrich. in Pharynx Expression | ParaSite Signal Peptide Prediction[e] | % of Residues in IDRs | Y. Kohara mRNA in situ expression pattern[f] | Additional Data Supporting Pharynx Expression (PMID)[g] | Mutant Phenotype in Pharnx (PMID) |
|---|---|---|---|---|---|---|---|---|---|---|---|
| 1 | abu-6 | APPG family (#1) member | old | C03A7.7 | 6, 263 | 72 | yes | 100 | yk309c1 | 15492775 (seq); 25361578 (transg) | |
| 2 | abu-7 | APPG family (#1) member | old | C03A7.8 | 6, 257 | 327 | yes | 100 | not tested | 15492775 (seq); 25361578 (transg) | |
| 3 | abu-8 | APPG family (#1) member | old | C03A7.14 | 6, 258 | 106 | yes | 100 | no pictures | 15492775 (seq) | |
| 4 | abu-10 | APPG family (#1) member | old | F35A5.3 | 6, 260 | 45 | yes | 100 | no pictures | 15492775 (seq); 28348061 (seq) | |
| 5 | abu-11 | APPG family (#1) member | old | T01D1.6 | 6, 240 | 35 | yes | 100 | no pictures | 15492775 (seq); 25361578 (transg); 21177967 (seq) | |
| 6 | abu-14 | APPG family (#1) member | old | ZK1067.7 | 5, 199 | 58 | yes | 100 | yk1393g06 | 15492775 (seq); 25361578 (transg); 15375261 (seq) | |
| 7 | abu-15 | APPG family (#1) member | old | C03A7.4 | 6, 262 | 144 | yes | 100 | no pictures | 15492775 (seq); 25361578 (transg) | |
| 8 | appg-1 | APPG family (#1) member | new | M02G9.1 | 4, 166 | 30 | yes | 100 | not tested | 17486083 (transg); 17850180 (transg); 20623595 (transg); | |
| 9 | appg-2 | APPG family (#1) member | new | F07H5.8 | 5, 206 | 52 | yes | 93 | yk32c6 | no published experiments found | |
| 10 | B0507.1 | no family assignment | old | B0507.1 | 4, 154 | 5 | yes | 0 | yk373f8 | 15492775 (seq); 18927627 (transg); 19372275 (transg) | |
| 11 | chs-2 | Chitin Synthase | old | F48A11.1 | 4, 152 | 49 | no | 25 | yk316g4 | 16098962 (transg) | |
| 12 | cht-1 | CHiTinase | old | C04F6.3 | 3, 116 | 12 | yes | 17 | yk109d2 (no pharynx) | no published experiments found | |
| 13 | cht-2 | CHiTinase | old | T13H5.3 | 4, 143 | 178 | yes | 5 | yk353g7 | no published experiments found | |
| 14 | cht-5 | CHiTinase | new | C08B6.4 | 3, 125 | 8 | yes | 0 | yk729c3 (no signal) | no published experiments found | |
| 15 | cht-6 | CHiTinase | new | T05H4.7 | 5, 184 | 30 | no | 0 | not tested | no published experiments found | |
| 16 | chtb-1 | CHiTin Binding | new | K04H4.2 | 3, 130 | 20 | yes | 0 | yk48g3 | 15492775 (seq) | |
| 17 | chtb-2 | CHiTin Binding | new | F23F12.8 | 4, 140 | 43 | yes | 59 | not tested | 17850180 (transg) | |
| 18 | chtb-3 | CHiTin Binding | new | T10E10.4 | 6, 235 | 49 | no, but likely SS | 0 | yk23a6 (no signal) | 15492775 (seq) | |
| 19 | chts-1 | CHiTosan Synthase | new | F48E3.8 | 4, 140 | 2 | no, but likely SS | 0 | yk272e1 (no signal) | 15492775 (seq) | |
| 20 | crm-1 | CRiM homolog | old | B0024.14 | 1, 24 | 6 | yes | 5 | yk42e12 (no pharynx) | 17486083 (transg); 17850180 (transg); 17869238 (transg) | |
| 21 | F30H5.3 | no family assignment | old | F30H5.3 | 3, 121 | 122 | yes | 0 | yk575d4 | 15492775 (seq) | |
| 22 | feh-1 | FE65 Homolog | old | Y54F10AM.2 | 6, 249 | 7 | no, but likely SS | 38 | yk23f9 | 11896189 (transg); 21177967 (seq) | this work |
| 23 | fipr-1 | FIP (Fungus-Induced | old | F23H12.8 | 8, 336 | 18 | yes | 61 | yk608e5 | no published experiments found | |
| 24 | fipr-2 | FIP Related | old | F23H12.9 | 8, 328 | 114 | yes | 0 | not tested | no published experiments found | |
| 25 | fipr-4 | FIP Related | old | C12D8.14 | 7, 297 | 101 | yes | 0 | not tested | no published experiments found | |
| 26 | fipr-6 | FIP Related | old | C12D8.17 | 7, 290 | 96 | yes | 0 | not tested | no published experiments found | |
| 27 | fipr-7 | FIP Related | old | C12D8.16 | 7, 312 | 70 | yes | 0 | not tested | no published experiments found | |
| 28 | fipr-9 | FIP Related | old | C12D8.19 | 7, 308 | 91 | yes | 0 | not tested | no published experiments found | |
| 29 | fipr-10 | FIP Related | old | C50H2.12 | 7, 311 | 24 | yes | 0 | not tested | no published experiments found | |
| 30 | fipr-11 | FIP Related | old | C50H2.10 | 7, 295 | 7 | yes | 0 | not tested | no published experiments found | |
| 31 | idpa-1 | IDP family A | new | T20B6.3 | 4, 173 | 46 | yes | 100 | not tested | no published experiments found | |
| 32 | idpa-2 | IDP family A | new | F07H5.6 | 5, 206 | 61 | yes | 82 | not tested | 15492775 (seq) | |
| 33 | idpa-3 | IDP family A | new | R07E3.2 | 5, 212 | 2 | yes | 91 | not tested | 15492775 (seq) | this work |
| 34 | idpb-1 | IDP family B | new | C04G6.10 | 5, 214 | 12 | yes | 100 | yk120a8 | 20623595 (transg) | |
| 35 | idpb-2 | IDP family B | new | C27A2.5 | 6, 226 | 7 | yes | 96 | not tested | 15492775 (seq) | |
| 36 | idpb-3 | IDP family B | new | C04G6.7 | 6, 228 | 30 | yes | 100 | no pictures | 21177967 (seq) | |
| 37 | idpb-4 | IDP family B | new | T01B7.8 | 6, 231 | 47 | yes | 100 | not tested | 15492775 (seq) | |
| 38 | idpb-5 | IDP family B | new | R13H4.8 | 6, 233 | 17 | yes | 74 | not tested | no published experiments found | |
| 39 | idpc-1 | IDP family C | new | Y47D3B.6 | 5, 215 | 105 | yes | - | yk529d4 | 21177967 (seq) | this work |
| 40 | idpc-2 | IDP family C | new | F41E6.11 | 6, 228 | 183 | yes | 100 | no pictures | no published experiments found | |
| 41 | idpc-3 | IDP family C | new | T06E4.12 | 6, 233 | 142 | yes | 100 | yk170a1 | no published experiments found | |
| 42 | idpc-4 | IDP family C | new | T06E4.14 | 6, 234 | 124 | yes | 100 | not tested | 21177967 (seq) | |
| 43 | idpc-5 | IDP family C | new | T06E4.8 | 6, 236 | 312 | yes | 88 | yk577d12 | 15492775 (seq) | |
| 44 | idpc-6 | IDP family C | new | T06E4.9 | 6, 240 | 164 | yes | 86 | not tested | 15492775 (seq) | |
| 45 | idpc-7 | IDP family C | new | T06E4.10 | 6, 253 | 116 | yes | 57 | not tested | no published experiments found | |
| 46 | idpp-1 | IDP enriched in Pharyx | new | T17H7.1 | 2, 72 | 145 | yes | 82 | yk815d04 | 17486083 (transg); 17850180 (transg) | |
| 47 | idpp-2 | IDP enriched in Pharyx | new | B0348.5 | 4, 138 | 7 | yes | 95 | not tested | no published experiments found | |
| 48 | idpp-3 | IDP enriched in Pharyx | new | F45B8.3 | 4, 168 | 386 | yes | 92 | yk663a1 | no published experiments found | |
| 49 | idpp-4 | IDP enriched in Pharyx | new | M02G9.2 | 5, 193 | 7 | yes | 85 | not tested | no published experiments found | |
| 50 | idpp-5 | IDP enriched in Pharyx | new | F35A5.4 | 5, 197 | 45 | yes | 100 | no pictures | 11997343 (transg) | |
| 51 | idpp-6 | IDP enriched in Pharyx | new | H22K11.3 | 5, 199 | 71 | yes | 100 | not tested | no published experiments found | |
| 52 | idpp-7 | IDP enriched in Pharyx | new | T23F6.1 | 5, 207 | 10 | yes | 87 | no pictures | no published experiments found | |
| 53 | idpp-8 | IDP enriched in Pharyx | new | M02G9.3 | 5, 217 | 42 | yes | 100 | not tested | no published experiments found | |
| 54 | idpp-9 | IDP enriched in Pharyx | new | T22B2.6 | 5, 221 | 17 | yes | 79 | not tested | no published experiments found | |
| 55 | idpp-10 | IDP enriched in Pharyx | new | D1007.13 | 6, 226 | 71 | no, but likely SS | 100 | not tested | no published experiments found | |
| 56 | idpp-11 | IDP enriched in Pharyx | new | C54D2.1 | 6, 245 | 227 | yes | 85 | not tested | no published experiments found | |
| 57 | idpp-12 | IDP enriched in Pharyx | new | W08E12.3 | 6, 249 | 19 | yes | 100 | yk720c3 | 30444224 (transg); 15492775 (seq); 17486083 (transg) | |
| 58 | idpp-13 | IDP enriched in Pharyx | new | F23H11.6 | 6, 254 | 20 | yes | 100 | not tested | no published experiments found | |
| 59 | idpp-14 | IDP enriched in Pharyx | new | C30F2.3 | 7, 307 | 37 | yes | 86 | yk253g10 | 28348061 (seq) | |
| 60 | idpp-15 | IDP enriched in Pharyx | new | ZC21.3 | 8, 339 | 49 | no, but likely SS | 100 | yk1099h07 | 7581455 (transg) | |
| 61 | idpp-16 | IDP enriched in Pharyx | new | F35B12.3 | 8, 357 | 68 | yes | 100 | not tested | no published experiments found | |
| 62 | itm-2 | InTegral Membrane protein | new | C25F6.7 | 3, 110 | 8 | no | 35 | yk82b10 | 28348061 (seq) | |
| 63 | lgx-1 | Lin-12 and Glp-1 X-hybrid. | old | C54G7.3 | 4, 151 | 26 | yes | 0 | yk208c10 | no published experiments found | |
| 64 | lrpc-1 | LRP10-like Chitin binder | new | M03E7.4 | 5, 188 | 54 | yes | 7 | not tested | 15492775 (seq) | this work |
| 65 | lrx-1 | LRP X(cross)-hybridizing | old | T04H1.6 | 4, 166 | 52 | yes | 32 | not tested | 15492775 (seq) | |
| 66 | myo-1 | MYOsin heavy chain | old | R06C7.10 | 8, 326 | 13 | | 0 | yk409g2 | 3320053 (Ab); 21177967 (seq) | 1603066 |
| 67 | myo-2 | MYOsin heavy chain | old | T18D3.4 | 7, 301 | 15 | no | 2 | yk127b7 | 3320053 (Ab); 7925019 (transg); 17486083 (transg) | 3320053 |
| 68 | myo-5 | MYOsin heavy chain | old | F58G4.1 | 7, 277 | 17 | no | 3 | yk310a4 | 17486083 (transg); 15492775 (seq); 17850180 (transg); | |
| 69 | nas-6 | Nematode AStacin protease | old | 4R79.1 | 4, 164 | 69 | yes | 0 | not tested | 20109220 (transg) | 20109220, this work |
| 70 | nep-1 | NEPrilysin metallopeptidase | old | ZK20.6 | 4, 143 | 9 | yes | 8 | yk4e1 | 16081104 (transg) | 16081104 |
| 71 | nep-12 | NEPrilysin metallopeptidase | old | F26G1.6 | 3, 134 | 21 | no | 12 | yk844g05 | 15492775 (seq) | |
| 72 | nspb-6 | NSPB family member | old | H04M03.2 | 7, 284 | 29 | yes | 0 | not tested | no published experiments found | |
| 73 | nspb-8 | NSPB family member | old | C01G12.10 | 7, 275 | 109 | yes | 0 | not tested | no published experiments found | |
| 74 | nspb-10 | NSPB family member | old | C01G12.6 | 7, 274 | 73 | yes | 0 | not tested | no published experiments found | |
| 75 | nspb-11 | NSPB family member | old | F35C5.10 | 7, 272 | 92 | yes | 0 | not tested | no published experiments found | |
| 76 | nspb-12 | NSPB family member | old | F09F7.8 | 7, 274 | 205 | yes | 0 | not tested | 15492775 (seq); 28348061 (seq) | |
| 77 | phat-4 | PHAryngeal gland Toxin-rel. | old | T05B4.3 | 5, 209 | 217 | yes | 0 | yk543d1 | 15492775 (seq) | |
| 78 | pqn-10 | APPG family (#1) member | old | C09E7.2 | 4, 160 | 85 | yes | 51 | not tested | no published experiments found | |
| 79 | pqn-13 | APPG family (#1) member | old | C14C11.8 | 4, 160 | 54 | yes | 98 | yk376c2 | 15492775 (seq) | |
| 80 | pqn-26 | APPG family member | old | DY3.5 | 4, 167 | 36 | yes | 86 | not tested | 15492775 (seq) | |
| 81 | pqn-29 | APPG family member | old | F10F2.9 | 5, 202 | 14 | yes | 86 | not tested | 15492775 (seq) | |
| 82 | pqn-31 | APPG family (#1) member | old | F21C10.8 | 8, 325 | 49 | yes | 100 | yk735h3 | 28348061 (seq) | |
| 83 | pqn-36 | APPG family (#1) member | old | F39D8.1 | 6, 260 | 62 | no, but likely SS | 98 | yk50g10 | no published experiments found | |
| 84 | pqn-54 | APPG family (#1) member | old | R09B5.5 | 6, 243 | 34 | yes | 97 | not tested | 15492775 (seq) | |
| 85 | pqn-63 | APPG family (#1) member | old | T06E4.11 | 5, 223 | 277 | yes | 92 | not tested | 15492775 (seq) | |
| 86 | pqn-71 | APPG family (#1) member | old | T23F1.6 | 7, 275 | 85 | yes | 100 | no pictures | 15492775 (seq); 28348061 (seq) | |
| 87 | pqn-74 | amyloid-chitin linker | old | W02A2.3 | 6, 232 | 115 | yes | 0 | yk506f12 | no published experiments found | |
| 88 | pqn-75 | APPG family (#1) member | old | W03D2.1 | 8, 345 | 29 | yes | 92 | yk520f8 | 28916707 (transg); 11997343 (transg) | 28916707 |
| 89 | pqn-94 | APPG family (#1) member | old | ZC15.8 | 8, 316 | 61 | yes | 70 | yk172g6 (ubiquit) | 21177967 (seq) | |
| 90 | R02F11.1 | no family assignment | old | R02F11.1 | 4, 160 | 48 | yes | 21 | yk597d1 | 15492775 (seq) | |
| 91 | tnc-2 | TropoNin C | old | ZK673.7 | 8, 337 | 9 | no | 0 | yk172g6 (no signal) | 15492775 (seq); 28508060 (seq) | 15743415 |
| 92 | tni-4 | TropoNin I | old | W03F8.1 | 8, 319 | 24 | no | 52 | yk328a10 | 15743415 (transg); 21177967 (seq) | 15743415 |
| 93 | tnt-4 | TropoNin T | old | T08B1.2 | 7, 307 | 18 | no | 66 | no pictures | 15492775 (seq) | |

**Supplemental Table 1. Genes of Special Interest and Evidence of Pharynx Expression.**

[a]All 78 gene products highlighted in Figure 4a are shown, along with 15 additional members of the *idpp* gene class referred to elsewhere in the text.

[b]The APPGs that have higher sequence similarity to one another and have more similar temporal expression patterns are described as APPG family (#1) members to distinguish them from more divergent APPGs.

[c]The Name Status indicates the 41 new WormBase-approved gene assignments.

[d]The indicated hour and degree is with respect to Figure 2a.

[e]In some cases, the updated Signal P algorithm will identify a signal peptide when ParaSite did not, as indicated with a 'no, but likley SS'.

[f]The spatial expression patterns of the indicated clones can be inspected at: http://nematode.lab.nig.ac.jp/dbest/srchbyclone.html . A green colour indicates confirmation of the expected expression pattern (enriched in pharynx); 'no signal' indicates little to no signal anywhere in photo micrographs. In two cases indicated in pink, signal could be observed in the animal, but the pharynx lacked signal.

[g]The PubMed ID number (PMID) is shown for the publication that provides additional spatial expression information for the gene. The nature of the data is either from a transgene (transg), an antibody (Ab), or is sequence-based (seq). A green colour indicates confirmation of the expected expression pattern (enriched in pharynx).

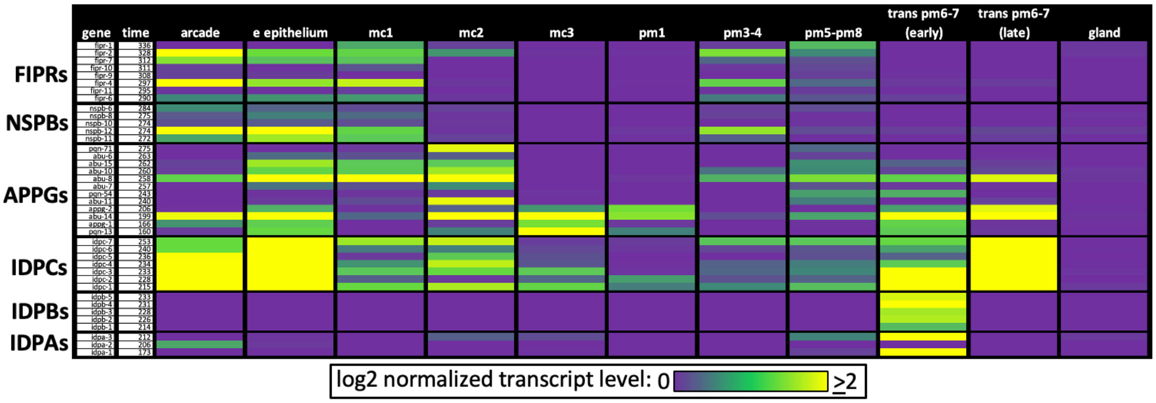


**Supplemental Table 2. Transcript Levels of Six Low Complexity Protein Families within Pharynx Cells.** The data is extracted from the data presented in Figure 9b.

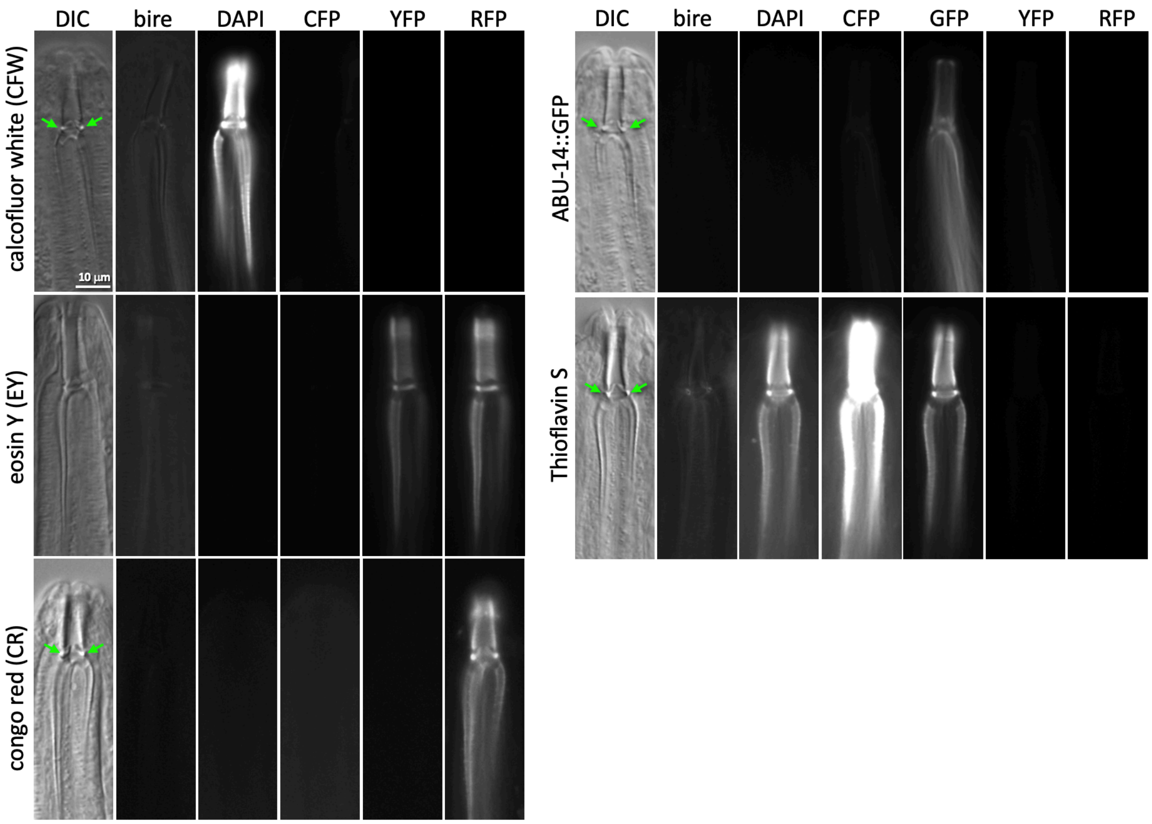


**Supplemental Figure 1. The Fluorescence and Filter Controls for Dye Staining.** Images of young adult wildtype worms incubated with the indicated dyes for 3 hours at the following concentrations: calcofluor white (CFW) (0.01% w/v), Eosin Y (EY) (0.15mg/mL), Congo Red (CR) (0.02% w/v), and ThS (0.01%). Also shown is an unstained ABU-14::GFP animal. Green arrows highlight the buccal collar dots that are often visible by DIC. The scale in A applies to all panels.

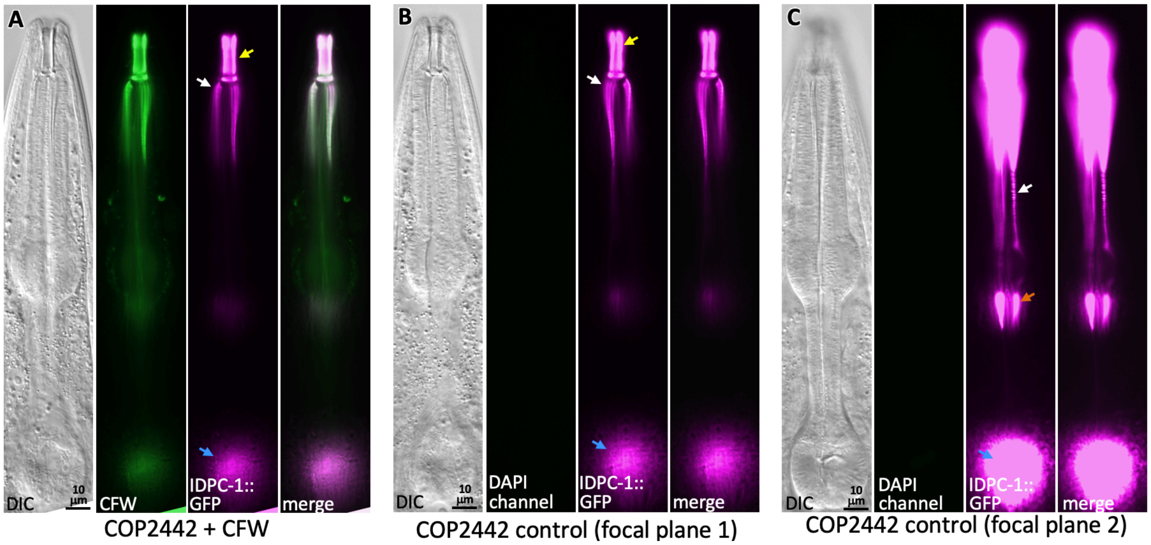


**Supplemental Figure 2. Expression Pattern of IDPC-1::GFP.** Shown is the *C. elegans* strain COP2442 *idpc-1::GFP(mGreenLantern)*. The GFP signal is specific to the pharynx and buccal cavity tissue. **A.** The same COP2442 animal depicted in Figure 5d but in a different focal plane that enables the visualization of the buccal cavity and channel cuticle. The animal is counterstained with the calcofluor white (CFW) chitin stain. **B&C.** A COP2442 animal not counterstained with CFW used as a control to show that the IDPC-1::GFP (green channel) signal does not bleed into the CFW (DAPI) channel. Two focal planes are shown. DIC, differential interference contrast; yellow arrows, the buccal cavity cuticle; white arrows, the corpus channels cuticle; blue arrows, the grinder of the terminal bulb; orange arrows, the isthmus channels cuticle; white arrows, the cuticle of the procorpus channels; yellow arrows, the cuticle of the buccal cavity.

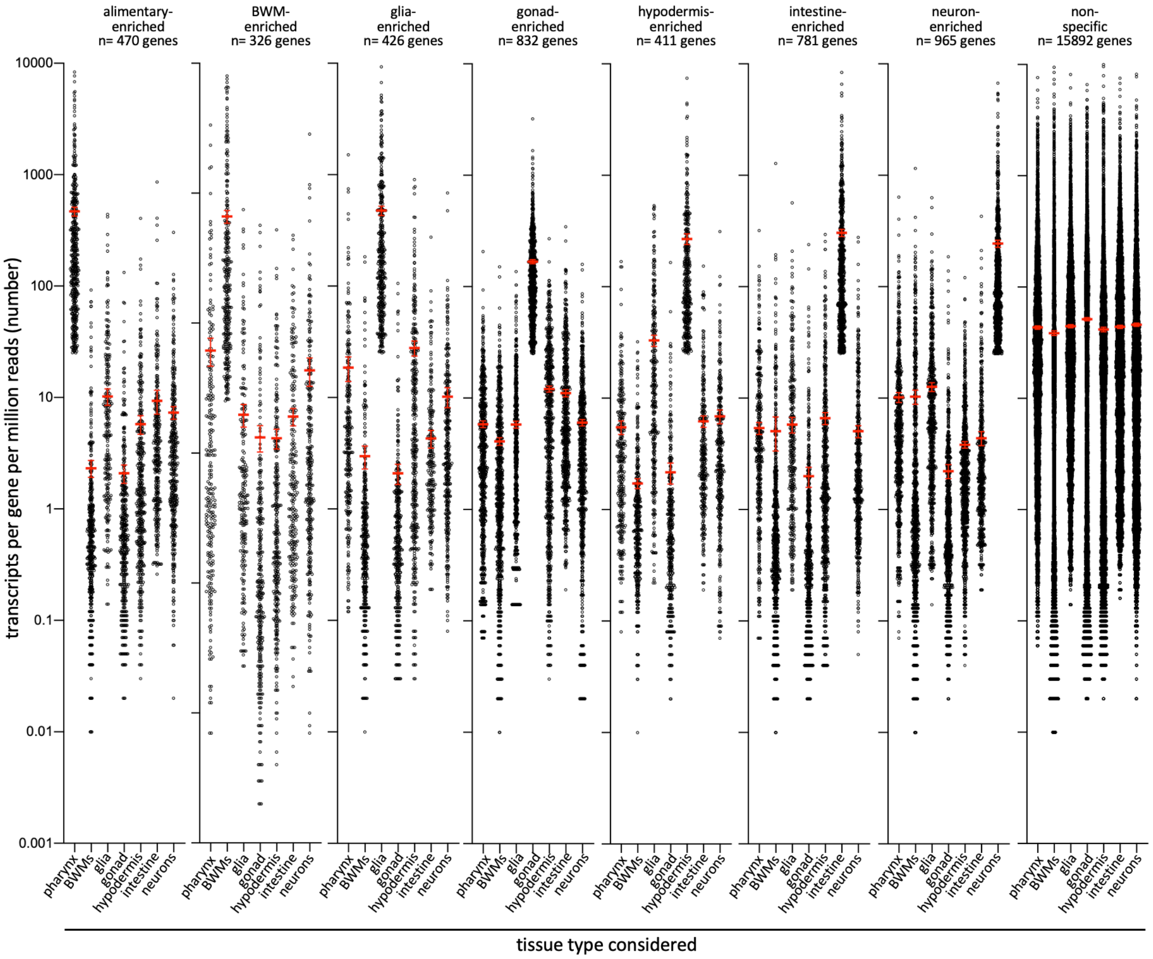


**Supplemental Figure 3. Tissue-Enriched Expression Levels of Tissue-Enriched Classes of Genes.** Each plot shows the expression level (as measured in single-cell sequencing analyses (*1*)) (y-axes) of the tissue-enriched set of genes in each tissue type (x-axis). Each dot represents the expression level for a single gene; each column reports the expression level of all genes in the respective set. For example, each of the 470 pharynx-enriched genes are expressed in the pharynx tissue at least 1.5 fold more than all other tissues combined. The plots demonstrate the enrichment of expression in the respective tissue type in each tissue-enriched gene set. The mean expression level for each tissue type and standard error of the mean is shown.

**A**

| secretome considered | number of query proteins | number of proteins in network | number of links | independent lines of evidence for links |
|---|---|---|---|---|
| pharynx | 261 | 20 | 36 | 209 |
| body wall muscles | 136 | 12 | 10 | 26 |
| glia | 228 | 0 | 0 | 0 |
| gonad | 36 | 0 | 0 | 0 |
| hypodermis | 214 | 0 | 0 | 0 |
| intestine | 262 | 31 | 56 | 119 |
| neurons | 319 | 3 | 3 | 10 |

**B**

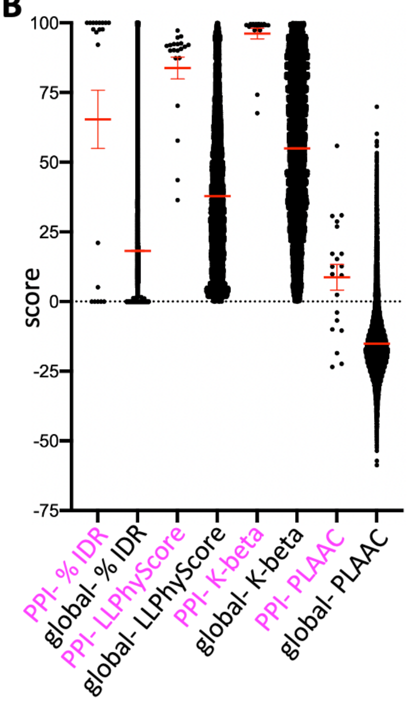


**Supplemental Figure 4. The Pharynx Secretome has a Greater Density of Protein-Protein Interactions (PPIs) Relative to other Secretomes. A.** The PPIs among gene products whose corresponding mRNAs are enriched in the indicated tissue (as derived from the L2 single-cell sequencing dataset (*1*)) and secreted (as defined by the presence of a signal sequence) (see Supplemental Data File 1 for details) as revealed through Genemania (*2*). Genemania settings used are 'equal by network weighting', and max resultant genes and attributes set to zero. Only networks that include clusters of 3 or more genes are considered here. Each interaction must be represented by multiple independent lines of experimental evidence. **B**. A comparison of the indicated properties between the group of 20 interacting proteins of the pharynx secretome and the entire proteome. All differences are significant ($p$<0.001) as measured by a Student's T-Test.

A. APPG consensus

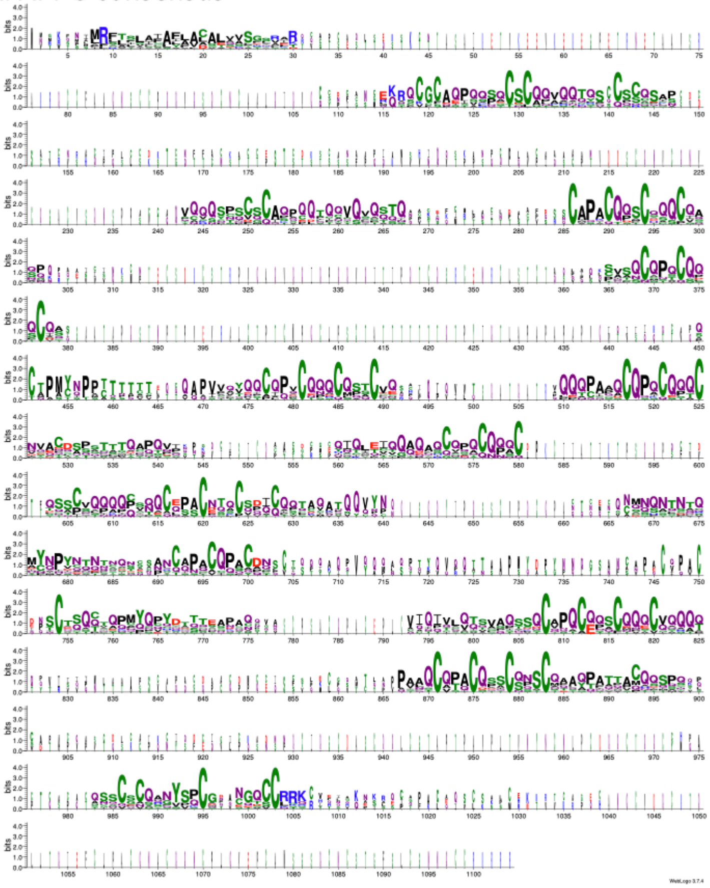

B. IDPA consensus

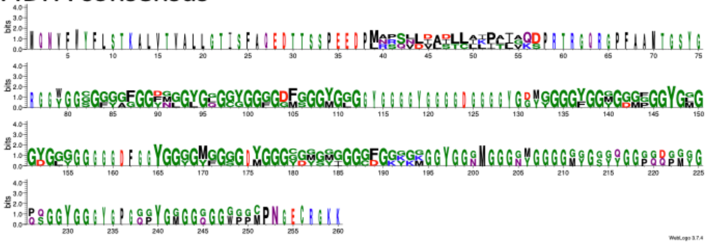

C. IDPB consensus

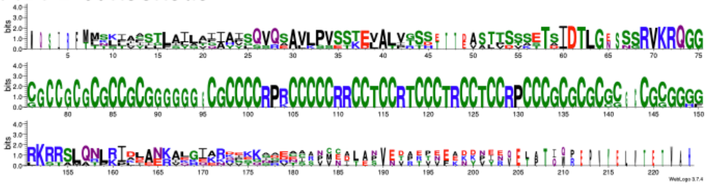

D. IDPC consensus

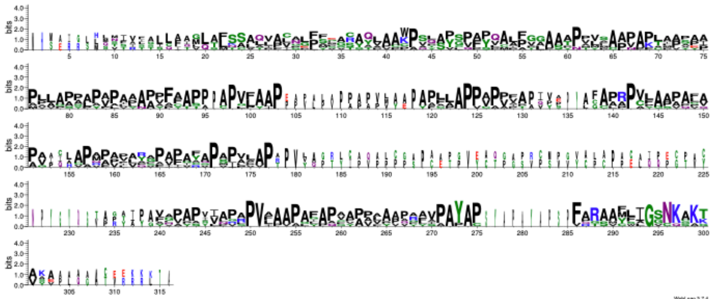

**Supplemental Figure 5. Consensus Sequence for Select Low Complexity Families.**

Sequence logos based on full length protein sequence alignments (*3*) of the APPGs (12 sequences), IDPAs (3 sequences), IDPBs (5 sequences) and IDPCs (7 sequences). Sequences were aligned with CLUSTALW and Sequence Logos were generated using WebLogo 3.7.4 (*4, 5*). Details of sequences used are found in Supplementary Data File 1.

**A**

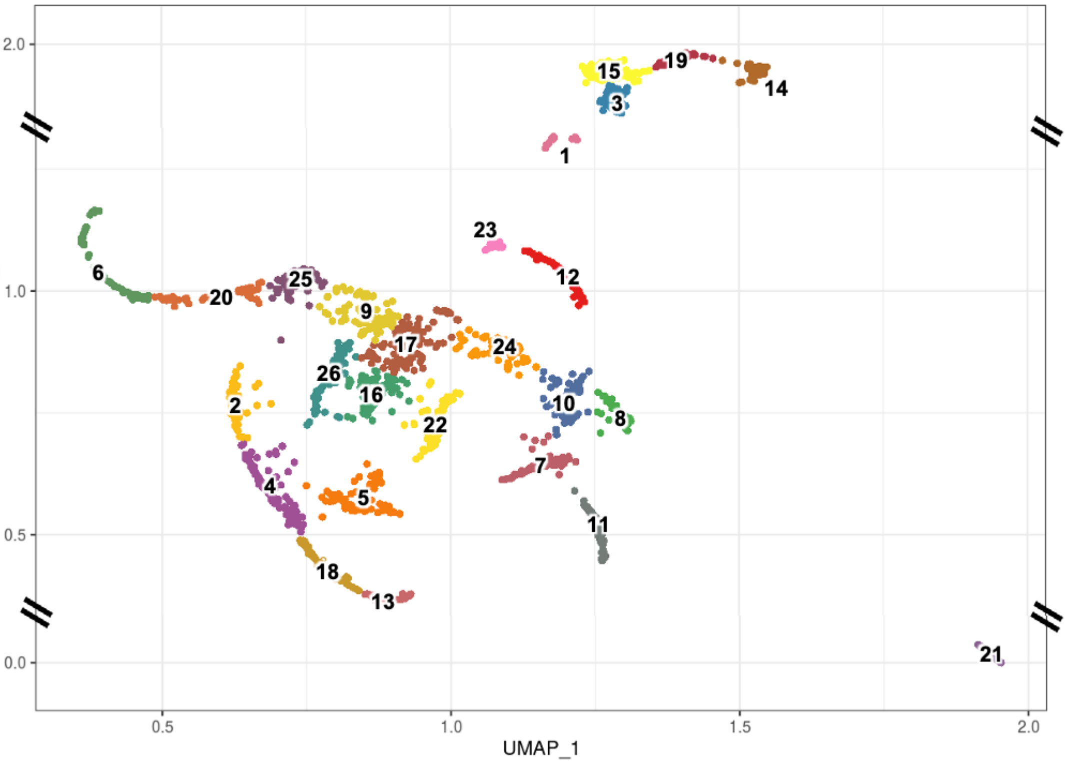


**B**

| Cluster | Cell Assignment | Evidence | Reference | PMID |
|---|---|---|---|---|
| 6, 20, 25 | aa and pa (arcades) | inx-5 | Altun et al 2009 | 19621339 |
| | | let-23 | van Buskirk and Sternberg, 2007 | 17891142 |
| | | hsp-43 | Sarov et al., 2012 | 22901814 |
| | | ajm-1 (epithelial gold standard) | Koppen et al., 2001 | 11715019 |
| | | no muscle gold stadards (myo-1, myo-2, mlc-1, mlc-2) | Rushforth et al., 1998 (mlc-1 and mlc-2), Okkema & Fire, 1994 (myo-2), Albertson, 1985 (myo-1 and myo-2) | 9799259 (mlc-1 and mlc-2), 7925019 (myo-2), 4054096 (myo-1 and myo-2) |
| 2, 4, 13, 18 | e epithelium | agr-1 | Hrus et al., 2007 | 17710131 |
| | | hsp-43 | Sarov et al., 2012 | 22901814 |
| | | sms-5, pgp-14 | Kamal et al 2019 | 31477732 |
| | | tat-3 | Lyssenko et al., 2008 | 18831765 |
| | | ifa-1 | Karabinos et al., 2003 | 14529618 |
| | | ajm-1 (epithelial gold standard) | Koppen et al., 2001 | 11715019 |
| | | no muscle gold stadards (myo-1, myo-2, mlc-1, mlc-2) | see above | see above |
| 26 | mc1 marginal | sms-5, pgp-14 | Kamal et al 2019 | 31477732 |
| | | tat-3 | Lyssenko et al., 2008 | 18831765 |
| | | ifa-1 | Karabinos et al., 2003 | 14529618 |
| | | marg-1 | Dineen & Gaudet, 2014 | 25480452 |
| | | inx-6 | Altun et al 2009 | 19621339 |
| | | ajm-1 (epithelial gold standard) | Koppen et al., 2001 | 11715019 |
| | | no muscle gold stadards (myo-1, myo-2, mlc-1, mlc-2) | see above | see above |
| 12, 23 | mc2 marginal | pgp-14 | Kamal et al 2019 | 31477732 |
| | | marg-1 | Dineen & Gaudet, 2014 | 25480452 |
| | | ifa-1 | Karabinos et al., 2003 | 14529618 |
| | | eat-5 | Altun et al 2009 | 19621339 |
| | | eat-5 | Starich et al, 1996 | 8707836 |
| | | glt-6 | Starich et al, 1996 | 8707836 |
| | | ajm-1 (epithelial gold standard) | Koppen et al., 2001 | 11715019 |
| | | no muscle gold stadards (myo-1, myo-2, mlc-1, mlc-2) | see above | see above |
| 7, 8, 10 | mc3 marginal (tentative) | nas-14 | Park et al., 2010 | 20109220 |
| | | nas-15 | Park et al., 2010 | 20109220 |
| | | ifa-1 | Karabinos et al., 2003 | 14529618 |
| | | no pgp-14 | Kamal et al 2019 | 31477732 |
| | | no muscle gold stadards (myo-1, myo-2, mlc-1, mlc-2) | see above | see above |
| 11 | pm1 | inx-1 | Altun et al 2009 | 19621339 |
| | | eyg-1 | Ian Hope, Don Moerman (personal communication via wormbase) | |
| unknown | pm2, pm4 | | | |
| 16 | pm3 | muscle gold stadards (myo-1, myo-2, mlc-1, mlc-2) | | |
| | | inx-10 | Altun et al 2009 | 19621339 |
| 22 | pm5-8 | muscle gold stadards (myo-1, myo-2, mlc-1, mlc-2) | see above | see above |
| | | inx-2, inx-3 | Altun et al 2009 | 19621339 |
| 3, 14, 15, 19, 21 | gland | lys-8, dod-6, hlh-6, B507.1, phat-1, phat-2, phat-4, phat-5, F41G3.10, phat-3 | Smit et al., 2008 | 18927627 |
| | | nas-5, nas-12 | Park et al., 2010 | 20109220 |
| 1, 5, 9, 17, | unknown | | | |

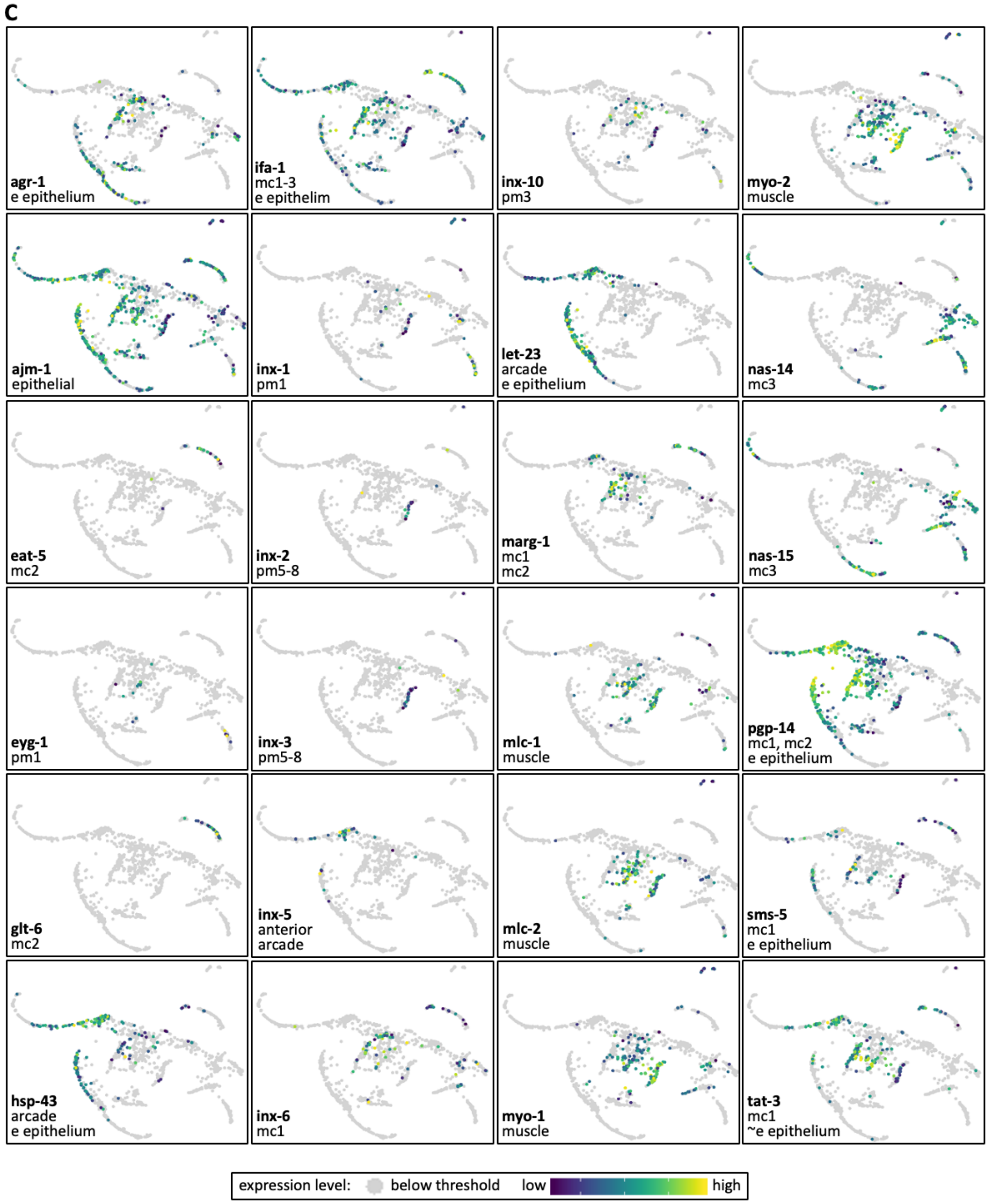


**Supplemental Figure 6. Identity Assignment of the Pharynx UMAP Clusters.**

**A.** The cluster map and cluster numbers are derived from Packer *et al* (*6*) using the single cell L2 sequencing data from Cao *et al* (*1*). Note the cropped Y axis for reasons of graphical clarity. **B.** Published gene expression patterns from transgenic reporters were used to assign identities to each of the clusters. Several of our assignments are different from that of Packet *et al* (*6*). **C.** The UMAP plots are extracted from the Packer *et al* dataset (see https://cello.shinyapps.io/celegans_L2/) and show the published markers used to assign identity to the clusters. Genes are arranged alphabetically and the cells in which GFP reporters serve as markers are indicated. For reasons of graphical clarity, the gland clusters are not shown.
